# Cryo-EM structures of engineered Shiga toxin-based immunogens capable of eliciting neutralizing antibodies with therapeutic potential against Hemolytic Uremic Syndrome

**DOI:** 10.1101/2024.12.21.626268

**Authors:** Alejandro Ezequiel Cristófalo, Arvind Sharma, María Laura Cerutti, Kedar Sharma, Roberto Melero, Romina Pardo, Fernando Alberto Goldbaum, Mario Borgnia, Vanesa Zylberman, Lisandro Horacio Otero

**Affiliations:** Centro de Rediseño e Ingeniería de Proteínas (CRIP), Escuela de Bio y Nanotecnologías (EByN), Universidad Nacional de San Martín (UNSAM), Consejo Nacional de Investigaciones Científicas y Técnicas (CONICET), 25 de Mayo 1401 (B1650HMQ), Gral. San Martín, Buenos Aires, Argentina; Genome Integrity and Structural Biology Laboratory, National Institute of Environmental Health Sciences (NIEHS), National Institute of Health (NIH), 111 TW Alexander Dr (NC 27709), Research Triangle Park, North Carolina, United States; Centro Nacional de Biotecnología (CNB), Consejo Superior de Investigaciones Científicas (CSIC), Calle Darwin 3 (28049), Madrid, Spain; Inmunova S.A, 25 de Mayo 1021 (B1650HMP), Gral. San Martín, Buenos Aires, Argentina; Departamento de Biología Molecular, Facultad de Ciencias Exactas, Físico-Químicas y Naturales, Instituto de Biotecnología Ambiental y Salud (INBIAS), Consejo Nacional de Investigaciones Científicas y Técnicas (CONICET), Universidad Nacional de Río Cuarto (UNRC), Ruta Nac. 36 - Km. 601 (X5804BYA), Río Cuarto, Córdoba, Argentina

**Author notes:** Corresponding author: Dr. Lisandro H. Otero, Departamento de Biología Molecular, Facultad de Ciencias Exactas, Físico-Químicas y Naturales, Instituto de Biotecnología Ambiental y Salud (INBIAS), Consejo Nacional de Investigaciones Científicas y Técnicas (CONICET), Universidad Nacional de Río Cuarto (UNRC), Ruta Nac. 36 - Km. 601 (X5804BYA), Río Cuarto, Córdoba, Argentina.

**Keywords:** Shiga-toxin-producing *Escherichia coli* (STEC), hemolytic uremic syndrome (HUS), Shiga toxin B subunits (Stx1B and Stx2B), *Brucella* spp. lumazine synthase (BLS), F(ab’)2 fragments, hyperimmune horse sera, chimera, protein structure, cryo-electron microscopy, antiserum, treatment, vaccine

## Abstract

Shiga toxin-producing *Escherichia coli*-associated hemolytic uremic syndrome (STEC-HUS) is a serious disease that causes renal failure predominantly in children. Despite its significant impact, there are currently no licensed vaccines or effective therapies available. The B subunits of Shiga toxins 1 and 2 (Stx1B and Stx2B) are suitable targets for developing neutralizing antibodies, but their pentameric assembly is unstable when isolated from the whole toxin. Taking advantage of the oligomeric symmetry shared between Stx1B and Stx2B with the lumazine synthase from *Brucella* spp. (BLS), we have previously engineered the chimeric toxoids BLS-Stx1B and BLS-Stx2B as immunogens to generate therapeutic equine polyclonal antibodies. The resulting product (INM004) has successfully passed phases 1 and 2 clinical trials, and a phase 3 has been launched in Argentina and seven European countries. In this work, we present the cryo-EM structures of BLS-Stx1B and BLS-Stx2B, which confirm that these engineered immunogens effectively stabilize the StxB pentamers. Moreover, our results reveal that both chimeric constructs present high flexibility at their extremes, corresponding to motions of the StxBs with respect to the BLS core. Additionally, we present structural evidence of the interaction between the chimeras and polyclonal Fab (pFab) fragments derived from INM004, demonstrating that the elicited neutralizing antibodies block most of the interaction surface of the toxins with their cellular receptors. These findings further validate this promising antibody-based therapy for mitigating STEC-HUS and demonstrate that the BLS-Stx1B and BLS-Stx2B chimeras are potential candidates for developing a human vaccine.

## INTRODUCTION

Shiga toxin-producing *Escherichia coli*-associated hemolytic uremic syndrome (STEC-HUS) is one of the leading causes of acute kidney injury (AKI) [1,2] in pediatric population and a relevant contributor to the development of chronic kidney disease (CKD) [3–5]. STEC-HUS is classified as an orphan pediatric disease with no licensed vaccines or specific treatment; therefore, only supportive therapy is currently advised for its clinical management. Children under five are the most impacted group [6], from which around 60% of patients require dialysis during the acute phase of the disease, with a mortality rate of approximately 3% [1,3]. In Argentina, STEC-HUS is endemic and has the highest incidence rates worldwide [7,8].

Infections caused by STEC are characterized by the production of Shiga toxins (Stxs), which are categorized into two main types—Stx1 and Stx2—and further subdivided into various subtypes. Among these, Stx2 is the most virulent, frequently associated with patients presenting the disease [9]. Despite the differences in toxicity, Stxs share a similar structural arrangement, in which a single A subunit (StxA) is assembled with five B subunits (StxB) (Fig. 1a) [10–15]. StxA is an enzymatically active 32 kDa monomer composed of two peptides—A1 (27.5 kDa) and A2 (4.5 kDa)—that inhibits protein synthesis in the affected cell. StxB is a homopentamer with a total molecular mass of 38.5 kDa (Fig. 1b), responsible for the internalization of the toxin by binding to the cell surface glycolipid globotriaosylceramide (Gb3) receptor.

**Figure 1.**
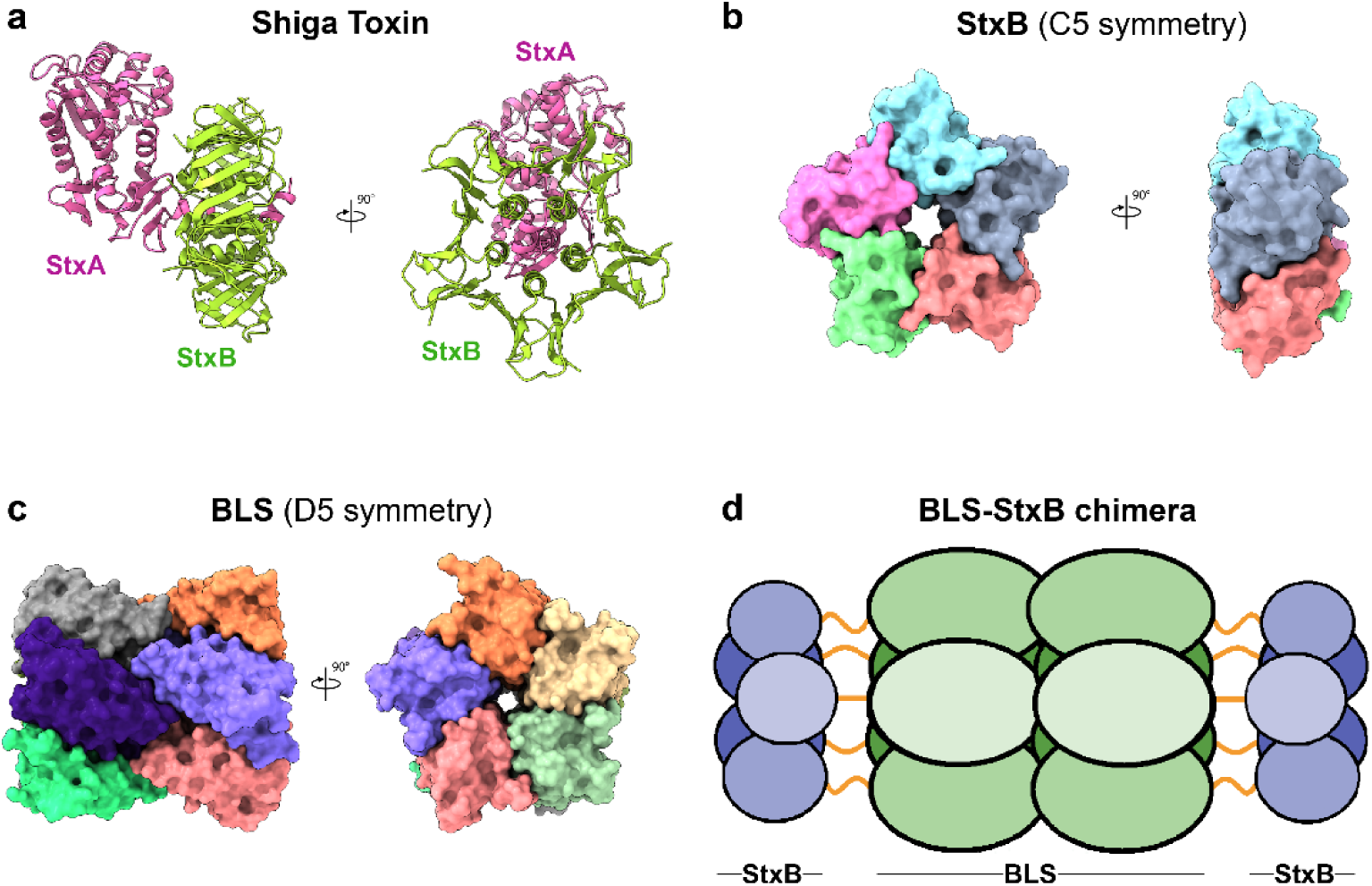
Oligomeric symmetry shared between StxB subunit and BLS. **a.** Crystallographic structure in ribbon representation of Stx2 (PDB code: 4M1U) with the StxA and StxB subunits colored in pink and green, respectively. **b.** Crystallographic structure in surface representation of the pentameric arrangement of Stx2B subunit, colored by chains (PDB code: 4M1U). **c.** Crystallographic structure in surface representation of BLS (PDB code: 1XN1) assembled as a dimer of homopentamers, colored by chains. **d.** Schematic representation of the postulated dipentameric model of BLS (green ellipses) engineered with Stx1B or Stx2B (blue circles) at the amino terminus. Linkers connecting both proteins are depicted as orange lines.

The role of Stxs makes STEC-HUS a toxemia rather than a systemic bacterial disease. Therefore, the neutralization of Stxs in compromised patients by passive immunization with antibodies offers a promising approach for treating the disease. In this context, the B subunit of Stx2 (Stx2B) emerges as a particularly attractive target for developing neutralizing antibodies, given its interaction with Gb3 and the high prevalence of Stx2 in patients with STEC-HUS. However, the potential of Stx2B as a target is limited by its poor immunogenicity [16], likely due to its unstable pentameric structure [17].

With the aim of stabilizing the oligomeric assembly of Stx2B, we devised a rational protein engineering strategy focused on the use of the lumazine synthase from *Brucella* spp. (BLS) as a structural scaffold. BLS is a highly immunogenic dimer of homopentamers with the versatility to display foreign antigens (Fig. 1c) [18,19]. By exploiting the shared oligomeric symmetry between the two proteins, we engineered Stx2B at the amino termini of BLS, creating a 254 kDa BLS-Stx2B chimera (Fig. 1d) [20]. This design aimed to preserve Stx2B in its native pentameric form, effectively mimicking the Gb3 binding sites of the full-length toxin.

The resulting chimera showed a remarkable capacity to elicit strong and long-lasting humoral immune responses against high lethal doses of Stx2 and its variants in mice, effectively inducing highly neutralizing antibodies [20–22]. Most importantly, sera from immunized mice also protected mice subjected to a relevant STEC infection model, demonstrating that the transferred antibodies could neutralize the toxin as it is delivered by the bacteria [20,21]. Prompted by these results, we also designed a second immunogen combining the B subunit of Stx1 (Stx1B) with BLS, resulting in the BLS-Stx1B chimera, with the aim of developing an antibody therapy with broader neutralizing capacity against different variants of Stxs [23].

Leveraging the strong immunogenicity of the BLS-Stx1B and BLS-Stx2B chimeras, we formulated a novel therapeutic product (hereafter referred to as INM004) comprising F(ab’)L fragments of equine polyclonal antibodies elicited by both immunogens [24]. This hyperimmune serum-based product demonstrated a very high neutralizing capacity against eight variants of Stx1 and Stx2 both *in vitro* and *in vivo*, potentially preventing the progression of STEC-HUS [23,25].

INM004 has obtained the Orphan Drug Designation in Europe by the European Medicines Agency (EMA) and in the United States by the Food and Drug Administration (FDA), where it also received the Rare Pediatric Disease Designation. Recently, phase 1 and 2 clinical trials were conducted for INM004 contributing to study the safety profile, tolerability, and pharmacokinetics of this medicine in a cohort of healthy adults and children having STEC-HUS [26,27]. No severe side-effects were detected during these studies, while only transient mild or moderate effects related to the infusion process were reported. Remarkably, INM004 retained its neutralizing capacity once infused in the bloodstream of the patients [26] and showed an adequate safety profile in pediatric STEC-HUS patients [27]. Furthermore, efficacy trends suggested a beneficial impact in mitigating kidney injury. These results encouraged progression to a phase 3 study of INM004 in pediatric patients with STEC-HUS [27], which has been recently launched in Argentina and nine European countries [28].

Looking ahead, INM004 has the potential to become the first targeted treatment for STEC-HUS patients worldwide. To further support the validation of this promising therapeutic product and explore possibilities for vaccine development, we present here the cryo-EM structural characterization of the BLS-Stx1B and BLS-Stx2B chimeras. Additionally, we provide structural evidence of their interactions with polyclonal Fab (pFab) fragments derived from INM004. These results provide perspective on the robust immunogenicity of the chimeric toxoids and the potent neutralizing capacity of the antibodies they elicit.

## RESULTS

### Cryo-EM structure determination of BLS-Stx1B and BLS-Stx2B chimeras

We conducted single-particle cryo-EM studies of both BLS-Stx1B and BLS-Stx2B chimeras to get atomic information on their molecular arrangements and validate the pentameric assembly of Stx1B and Stx2B when engineered with BLS. Details on data acquisition, data processing, and map statistics are described in the Materials and Methods section, Table 1, and Fig. S1 and S2.

**Table 1.**
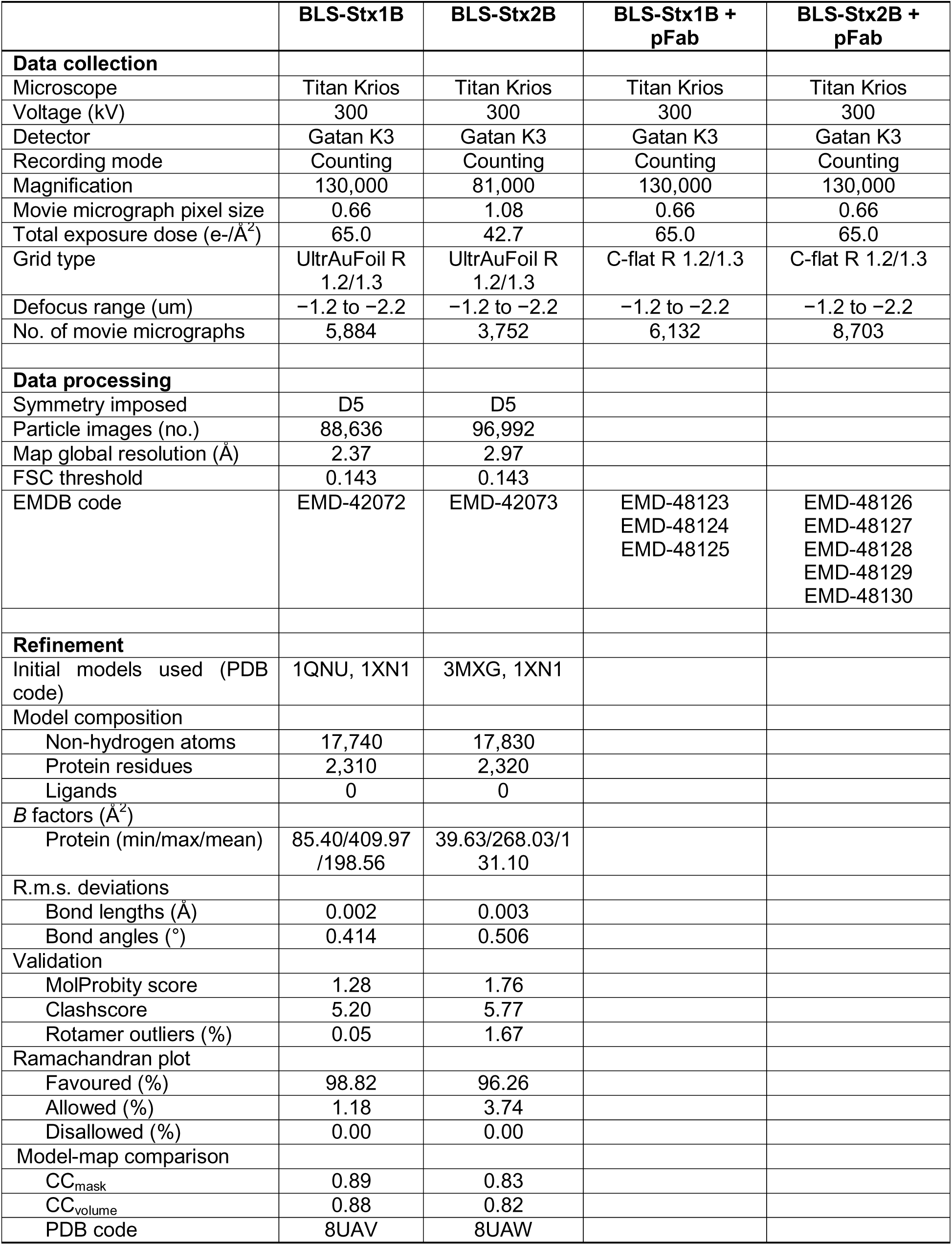
Cryo-EM data collection, refinement, and validation statistics.

The examination of the 2D classes of both proteins clearly unveiled side and top views of the particles resembling the proposed model, with the StxB subunits at both extremes of the BLS structure (Fig. 2a). The 3D reconstructions with imposed D5 symmetry yielded overall resolution maps of ∼2.4 Å for BLS-Stx1B and ∼3.0 Å for BLS-Stx2B, with a maximum diameter of ∼150 Å (side view) and a minimum diameter of ∼80 Å (top view) (Fig. 2b). Remarkably, both maps revealed higher details for BLS than for the StxB subunits. This is also evidenced by the local resolution of the maps, which was around 2.4–3.0 Å for BLS and 6.0–7.0 Å for the StxBs (Fig. 2c).

**Figure 2.**
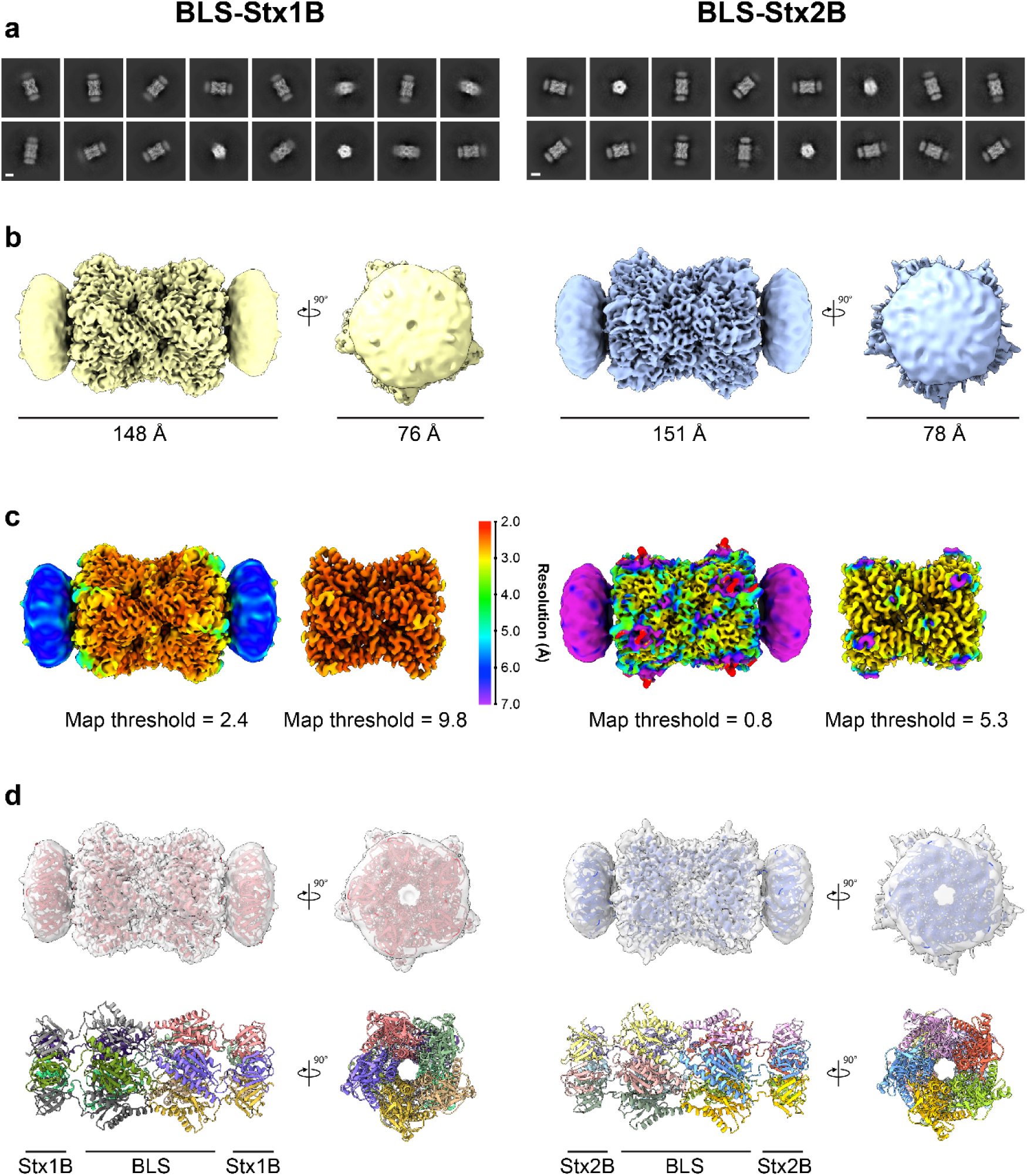
Cryo-EM structures of BLS-Stx1B (*left*) and BLS-Stx2B (*right*) chimeras. **a.** Representative reference-free 2D class averages with a 50 Å scale bar. **b.** Final 3D maps in two orientations (side view and top view) colored by estimated local resolution. Scale bars are displayed for reference. **c.** Local resolution maps at two map thresholds. **d.** *Upper*: Atomic models fitted into the density maps. The models are shown in ribbon representation colored in red (BLS-Stx1B) and blue (BLS-Stx2B) and displayed with the maps (gray) transparently overlaid. *Bottom*: Atomic models shown in ribbon representation, colored by chains.

Based on the obtained density maps, we built atomic models for both chimeras (Table 1, Fig. 2d, and Fig. S3). The regions of BLS and the StxB subunits were modeled based on previously described crystallographic structures, as described in the Materials and Methods section, while the linker regions were manually built from the ground up. In both chimeras, the structure of BLS was identical to the previously reported crystallographic model (PDB: 1XN1) [29,30] (Cα-r.m.s.d. values of 0.596 Å and 0.662 Å for BLS-Stx1B and BLS-Stx2B, respectively), showing a decameric assembly formed by a 17 kDa subunit arranged as a dimer of pentamers (Fig. 2d, and Fig. S3). On the other hand, although the regions of the toxins in the 3D maps were not defined in detail, the crystallographic models of both StxB pentamers (Stx1B, PDB: 1QNU [15] and Stx2B, PDB: 3MXG [29]) properly fitted into them (Fig. 2d, *upper,* and Fig. S3). According to the models, the pentamers of both proteins align to form a central pore, giving the chimeric structure a D5 symmetry (Fig. 2d, *bottom*). This symmetry compatibility between the quaternary structures of BLS and the StxB subunits, essential for the rational design of the engineered immunogens outlined here, is further confirmed by our structural data.

The low resolution observed in the Stx1B and Stx2B suggests a high degree of motion when these subunits are attached to the BLS scaffold, likely facilitated by the highly flexible linkers. Indeed, the continuous heterogeneity observed precluded 3D classifications that could otherwise yield higher-resolution maps of their distinct conformations. Prompted by these results, we decided to deepen the study on the movement of the StxB subunits by applying 3DFlex from CryoSPARC, a recently described motion-based neural network model for continuous molecular heterogeneity [31]. Following the workflow suggested by the developers, we started preparing the parametrized meshes that depict the rigid and flexible regions of the proteins using the previously obtained consensus 3D maps of the chimeras. We then trained the corresponding motion models, ultimately obtaining four latent coordinates for each chimera, which captured in greater detail the distinct movements of the StxB subunits with respect to the BLS scaffold (Fig. 3 and Supplementary movies 1-4 (BLS-Stx1B) and 5-8 (BLS-Stx2B)). These structural fluctuations impart substantial flexibility to the chimeric constructs, particularly at their extremes, resulting in overall non-rigid structures. Importantly, despite this significant flexibility, the StxB assemblies preserve their pentameric conformation upon association with the BLS scaffold, underscoring the structural robustness of the chimeric constructs.

**Figure 3.**
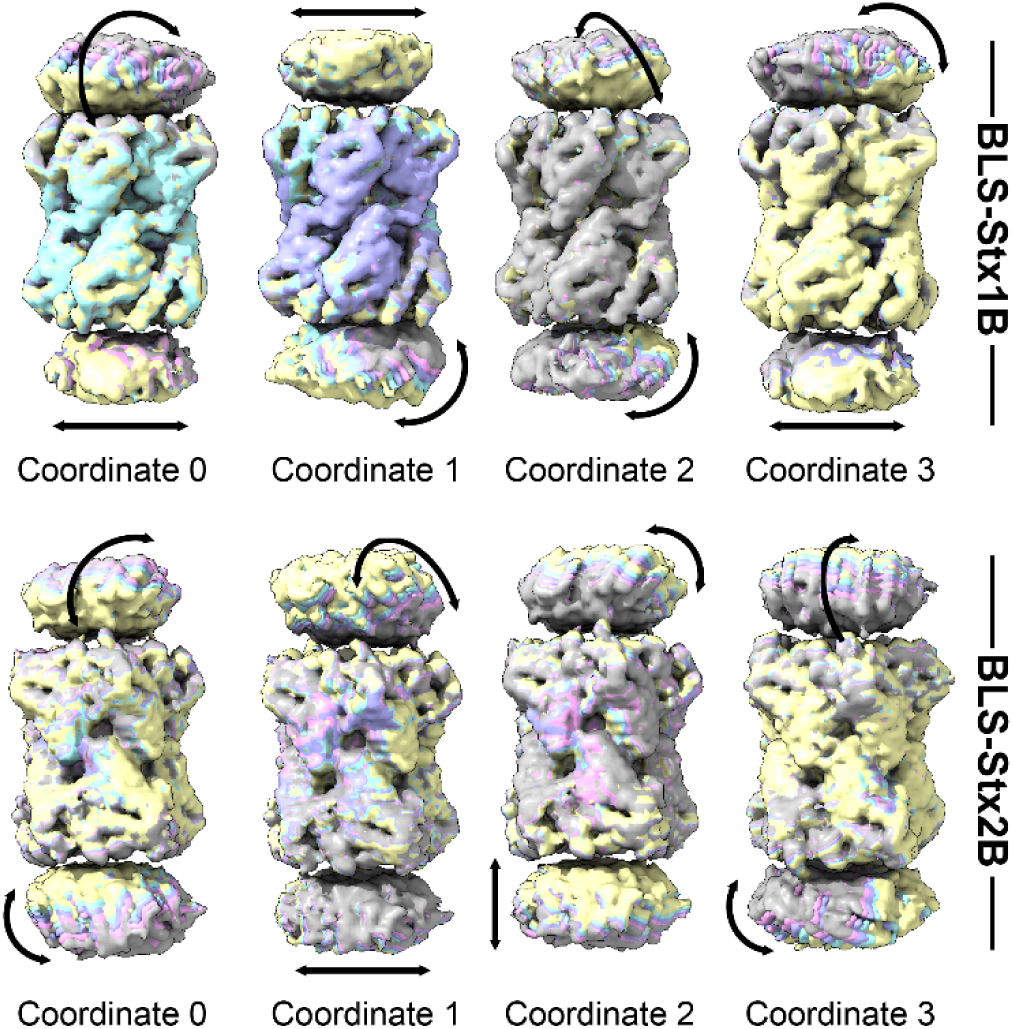
Continuous molecular heterogeneity obtained by 3DFlex for BLS-Stx1B (*upper*) and BLS-Stx2B (*bottom*) chimeras. Colored series of convected densities at five positions are depicted for each of the four latent coordinates learned by 3DFlex in both chimeras. Each dimension shows a different type of motion within the same model. The arrows represent the mode of movement of the toxin pentamers for each coordinate. For details see Supplementary movies 1-4 (BLS-Stx1B) and 5-8 (BLS-Stx2B)

### Cryo-EM analysis of the immune complexes between the chimeras and the pFab fragments derived from INM004

We aimed to visualize and characterize the complexes formed between the BLS-Stx1B and BLS-Stx2B immunogens and the induced neutralizing antibodies that constitute the therapeutic product INM004.

A method to study the polyclonal response to different antigens performing a structural epitope mapping through cryo-EM was previously reported by Ward et al. and applied to different systems [32–35]. In their works, the whole starting sera from patients and/or immunized animals were digested and incubated with the corresponding antigens to obtain Fab-bound complexes that were SEC-purified prior to EM grid preparation.

Given that INM004 contains both specific (anti-Stx1B and anti-Stx2B) and non-specific immunoglobulins, in our approach, the F(ab’)_2_ fractions present in INM004 were purified through three sequential affinity chromatography steps (see Materials and Methods section for details). First, we used a BLS-activated column to deplete the sample from anti-BLS antibodies, which are not therapeutically relevant. Next, we performed successive purification steps using BLS-Stx1B and BLS-Stx2B activated columns to obtain separated pools of specific anti-Stx1B and anti-Stx2B antibodies. The corresponding anti-Stx1B and anti-Stx2B F(ab’)_2_ fragments were then digested to obtain specific pFab fragments, which were purified and further incubated with their corresponding chimeras. Prior to cryo-EM examination, the complexes were SEC-purified to separate them from unbound pFabs. This approach enabled us to obtain the specific populations of the complexes formed between BLS-Stx1B and BLS-Stx2B with their respective anti-Stx1B and anti-Stx2B pFabs, simplifying the samples for EM imaging and data processing and analysis.

Vitrified samples of the complexes were imaged by cryo-EM (Table 1), revealing predominantly smaller-than-expected particles, which resulted in micrographs with a largely noisy background (Fig. S4 and S5). Despite several attempts to optimize the sample vitrification, the presence of these particles remained consistent. We hypothesized that they could correspond to unbound Fabs, likely stemming from the partial disruption of the antigen/antibody complexes during sample manipulation and/or freezing. For the image processing, we first used CryoSPARC Blob picker [36] adjusting the blob size according to the diameter of the chimeras’ models considering an extension due to the bound pFabs. After conducting 2D classification, our primary findings revealed predominant classes that roughly resembled top views of the pentameric models but, in contrast to observations of the uncomplexed chimeras, the characteristic elongated side views were conspicuously absent, preventing the corresponding full-length 3D reconstructions.

In an attempt to mitigate the preferential orientation bias towards top-like distribution and to reconstruct the complete structure of the complexes, we used Topaz [37] models trained with BLS-Stx1B and BLS-Stx2B particles obtained from the previously described data and performed Topaz extract jobs in CryoSPARC. After several 2D classification rounds of the selected particles, we detected 2D classes representing different views of the complexes, including side views (Figs. S4 and S5). These particles were subsequently used to generate single-class 3D maps with no symmetry imposed of the BLS-Stx1B/pFabs and BLS-Stx2B/pFabs complexes at ∼8 Å resolution(Figs. S4 and S5, *insets*). The volumes clearly revealed additional densities in the region of one of the two Stx pentamers, consistent with the presence of bound Fabs. The absence of extra density on the opposite pentamer could not be determined, though we speculate it may be attributed to the predominant top-like orientation present in the data.

In an effort to improve the maps and gain details of the pFab-bound complexes, we conducted additional data processing as follows. The corresponding particles were 3D reconstructed imposing D5 symmetry using a low-pass filtered volume of the BLS region as initial reference and a mask enclosing this part of the volume. Then, the particles were symmetry expanded and subjected to focused 3D classification using focalized masks over each observed extra density. After 3D reconstruction of the expanded particles, we obtained maps of individualized complexes for both chimeras (Figs. S4 and S5). In both cases, crystallographic Fab models were fitted into the additional density regions, resulting in the identification of three and five distinct complexes for BLS-Stx1B and BLS-Stx2B, respectively (Figs. 4a and 4b). Interestingly, both complexes display distinct orientations and positions of the pFabs, indicating differences in the recognized epitopes (Fig. 4c).

**Figure 4.**
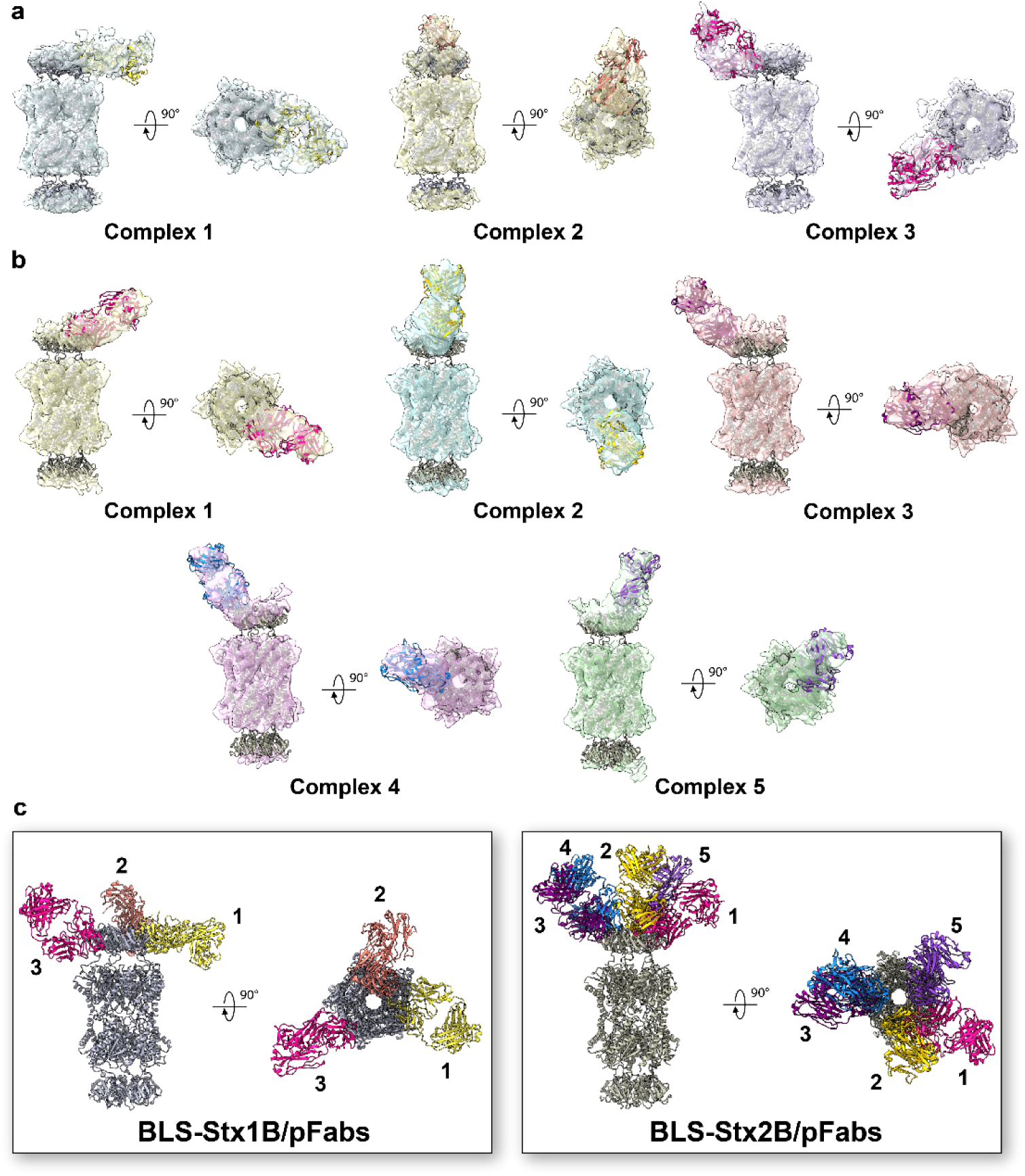
Cryo-EM structures of the complexes formed between the chimeras and the pFab fragments. a-b. Atomic models fitted into the individualized 3D maps for BLS-Stx1B/pFabs (a) and BLS-Stx2B/pFabs (b) complexes shown in two orientations (side view and top view). Atomic models of the chimeras and the Fab fragments are displayed in different colors with the maps transparently overlaid. **c.** Combined atomic models shown in ribbon representation of BLS-Stx1B/pFabs (*left*) and BLS-Stx2B/pFabs (*right*) complexes. The chimeras are shown in gray, with the Fabs fragments labeled by numbers and distinguished by various colors.

To assess whether the epitopes recognized by the different pFabs overlap with the cellular receptor-binding sites of Stxs, the crystallographic models of Stx1B (PDB: 1BOS) [14] and Stx2B (PDB: 2BOS) [38] in complex with Gb3 oligosaccharides were aligned to the corresponding pFab-bound chimera (Fig. 5). Notably, the defined pFabs extensively cover most of the Gb3-binding sites on both StxBs. In the case of BLS-Stx2B, the elicited antibodies appear to recognize and hide a larger surface area compared to BLS-Stx1B. In both cases, this structural analysis supports the potent and broad neutralizing capacity of INM004.

**Figure 5.**
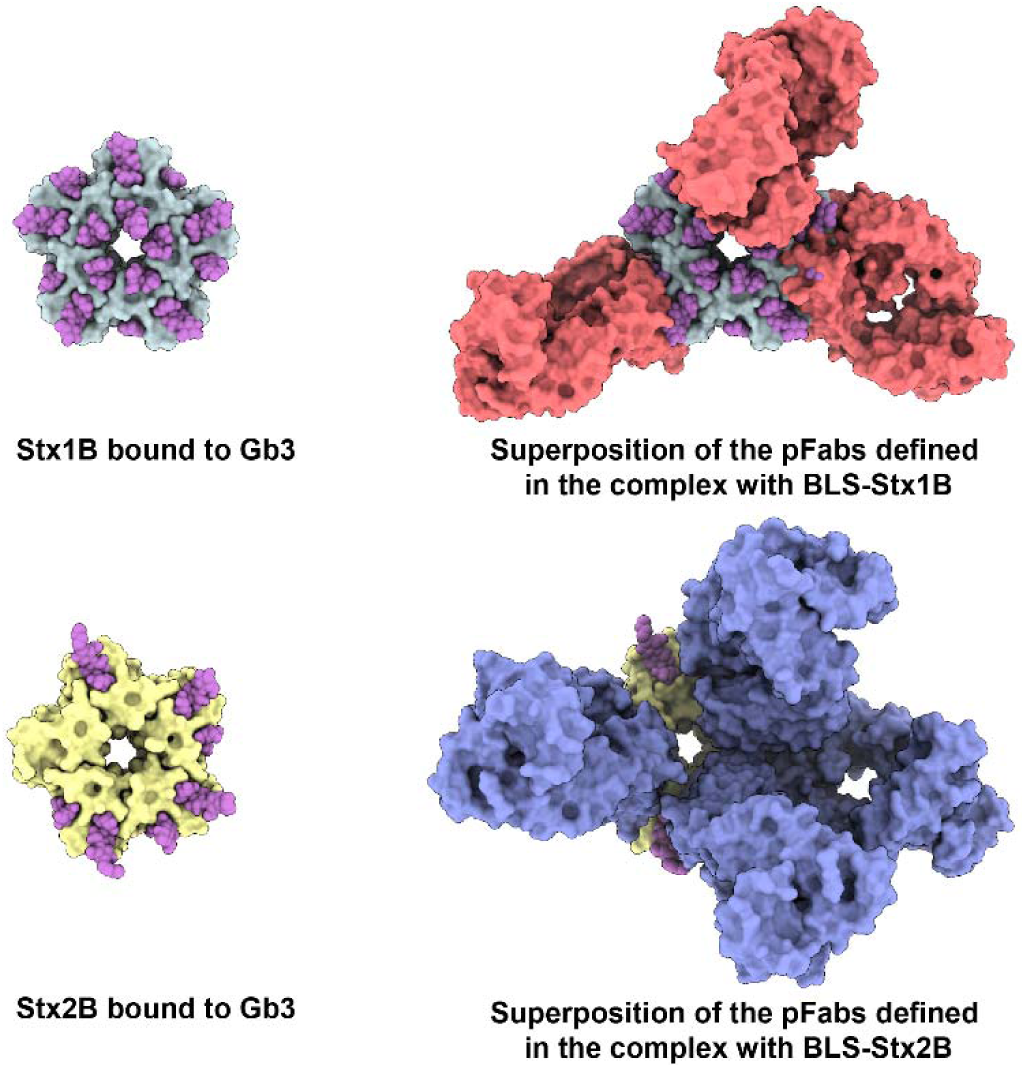
Structural comparison of the receptor-binding sites of Stx1B and Stx2B with the epitopes recognized by the pFabs derived from INM004. Top view of the crystallographic models displayed as surface representation of Stx1B in grey (PDB: 1BOS) and Stx2B in yellow (PDB: 2BOS), both bound to Gb3 oligosaccharides, depicted as magenta spheres (*left*). Structural superposition of the crystallographic StxB/Gb3 complexes with the atomic coordinates of the pFab models presented in Fig. 4 (*right*). The models of the chimeras are hidden for clarity, while the pFab fragments are shown as surface representation in red (complex with BLS-Stx1B) and blue (complex with BLS-Stx2B).

## DISCUSSION

Protein assemblies with a high degree of repetitiveness and organization are known to induce strong immune responses [39]. This is why polymeric display of antigens is a powerful protein engineering strategy that enhances the design of immunogens, leading to the development of vaccines and antibody-based therapies [40]. However, displaying whole proteins onto oligomeric scaffolds can be constrained by an unsuitable structure of the foreign protein and its propensity to undergo homomeric interactions.

To the best of our knowledge, there are only a few successful cases of polymeric protein scaffolds used to display other polymeric proteins. A notable example is the application of hepatitis B virus capsid-like particles for displaying the dimeric outer surface lipoprotein C (OspC) from *Borrelia burgdorferi*, the causative agent of Lyme disease [41]. However, it is important to note that most of these cases lack a thorough polydispersity analysis.

Leveraging the shared oligomeric symmetry between Stx1B and Stx2B with the highly immunogenic BLS enzyme, we previously engineered the chimeric toxoids BLS-Stx1B and BLS-Stx2B as immunogens to develop therapeutic polyclonal antibodies [20,21,26,27] and monoclonal neutralizing antibodies [22]. Nonetheless, this approach faced significant challenges [24]. First, the polymerization of the StxBs could occur between different protein particles, resulting in cross-linking and massive aggregation. Second, the kinetics of folding and association of the StxBs and BLS pentamers could differ significantly, potentially trapping the fusion particle in an intermediate folding state. Third, the structural constraints imposed by the covalent link to the pentamers of BLS could not be adequate to stabilize the StxB pentameric structure. Fourth, the topology of the oligomeric scaffold could not be compatible with that of the oligomeric target. Fifth, the interactions between intermediate configurations of both proteins could prevent the proper folding and assembly of the fusion particles. Sixth, the low stability or low solubility of the StxBs could adversely affect the stability of the whole particles inducing aggregation of the chimeras.

The experimental structures of BLS-Stx1B and BLS-Stx2B presented here validate the previously proposed models dispelling most of the potential issues mentioned above and explain, at near-atomic resolution, the reasons behind the enhanced immunogenicity of these chimeras.

In contrast to the tendency of isolated StxBs to dissociate their pentameric assemblies, they remain associated in the chimeric structures. The stable oligomeric scaffold of BLS provides a supportive framework, effectively substituting the role of the α-helix from StxA, which naturally inserts into the center of the pentamer in native toxins [12]. However, this BLS-assisted chimeric assembly is not rigid; our results demonstrate that the StxBs can move significantly as a whole due to the flexibility of the linkers connecting the two proteins. This highlights the dynamic behavior of the regions involved in eliciting neutralizing antibodies within the chimeras.

Beyond the inherent heterogeneity of polyclonal antibodies, which makes structural studies challenging, obtaining information on their modes of interaction is crucial for optimizing the corresponding immunogens. This is especially important in applied contexts such as the development of vaccines and therapeutic antibodies, where precise structural insights can significantly enhance the efficacy and safety of the resulting products. In the immune complexes presented in this study, the challenges associated with antibody heterogeneity were further exacerbated by the high flexibility of the chimeras and the limited number of particles, which made imaging and processing even more difficult. Nevertheless, the results obtained clearly demonstrated that the pFabs from INM004 effectively cover most of the interaction surface of the toxins with their cellular receptors. In addition, we evidenced that the elicited pFabs interact distinctly with both chimeras, suggesting differences in epitope accessibility between Stx1B and Stx2B. Since INM004 can strongly neutralize the eight clinically relevant Stxs [23], this experimental approach helps to explain the structural reasons for its efficacy.

These findings provide valuable insights into the mechanism of action behind this promising antibody-based therapy for mitigating STEC-HUS, which is currently being evaluated in a phase 3 clinical trial [28]. They not only enhance our understanding of how the therapy works but also underscore its potential as an effective approach for reducing the impact of the disease. Furthermore, it can be hypothesized that children immunized with a mixture of these chimeras would generate neutralizing antibodies against all clinically relevant Stxs, further supporting their potential use as a human vaccine against STEC-HUS.

Finally, this study also contributes to highlighting the capability of cryo-EM to capture dynamic molecular complexes, such as those involved in antigen–antibody interactions. This makes it a particularly useful tool for mapping epitopes that induce protective immune responses, enabling the optimization in the design of immunogens for more effective vaccines and antibody-based therapies.

## MATERIALS AND METHODS

### Protein expression and purification

Expression and purification of BLS-Stx1B [23] and BLS-Stx2B [20] were performed as previously described. Briefly, pET11a vectors containing the gene sequence of the corresponding chimera were transformed into competent *E. coli* BL21 (DE3) cells. Inclusion bodies containing most of the protein were solubilized overnight in 8 M urea, 50 mM Tris-HCl, 5 mM EDTA, pH 8.0. The unfolded material was purified by ionic exchange chromatography in a Q-sepharose column using a Dionex Ultimate 3000 HPLC apparatus. The elution was performed using a linear gradient from 0 to 1 M NaCl in 8 M urea, 50 mM Tris-HCl, pH 8.5. Subsequently, the chimeras were refolded by dialysis against a buffer containing 10 mM NaLHPO, 1.8 mM KHLPO, 137 mM NaCl, 2.7 mM KCl, pH 7.0, followed by centrifugation at 18,000 g for 20 minutes. The soluble fraction was then collected and filtered through a 0.22 µm filter. Recombinant BLS used for activating the affinity chromatography column was expressed in *E. coli* BL21 (DE3) cells carrying the pET11b plasmid and purified thrugh ion exchange chromatography, as previously described [42].

### Preparation of pFab fragments

The preparation of INM004, consisting of polyclonal F(ab’)_2_ fragments obtained from sera of horses immunized with BLS-Stx1B and BLS-Stx2B, was performed as described previously [23]. INM004 was subjected to a further serial purification procedure through affinity chromatography to obtain specific F(ab’)_2_ fragments against Stx1B and Stx2B. To that aim, three HiTrap NHS-activated affinity columns (Cytiva) were pre-activated with BLS, BLS-Stx1B, and BLS-Stx2B following the instructions provided by the manufacturer. First, INM004 was passed through the BLS-activated column to deplete the sample of anti-BLS fragments. The sample was introduced in the column and after one minute incubation at room temperature, binding buffer (20 mM PBS + 0.15 M NaCl, pH 7.4) was used to elute the unspecific fragments. Elution buffer (100 mM glycine + 0.5 M NaCl, pH 3.0) was used to elute the specific fragments, which were immediately neutralized with a buffer containing 1 M Tris-HCl, pH 8.8. The elutions were controlled by Bradford colorimetric assay and subsequent absorbance reading at 280 nm. The flow-through (FT) of this chromatography was used as input for the next purification step using the BLS-Stx1B-activated column following the same procedure. The fraction containing the anti-Stx1B pFabs was preserved. Finally, the FT of the second chromatography was used as input of the BLS-Stx2B activated column, which was eluted as described above.

The fractions of the specific F(ab’)_2_ fragments were pooled, concentrated and buffer-exchanged to digestion buffer (1X PBS + 10 mM EDTA + 12.5 mM β-mercaptoethanol, pH 7.0) in Amicon 50 kDa MWCO ultra centrifugal filters (Millipore). Later, F(ab’)_2_ samples were treated with papain-agarose resin (Thermo Fisher Scientific) at 37 °C for 4 h using a 1:200 p/p enzyme:F(ab’)_2_ ratio. The obtained pFab fragments were further purified through Size-Exclusion Chromatography (SEC) in a Superdex200 increase column (Cytiva). Chromatography was performed at a flow rate of 0.5 mL/min using a Waters 600E Multisolvent Delivery System. The recorded signal was set at 280 nm.

### Cryo-EM sample preparation

For the preparation of the chimeras, samples of BLS-Stx1B and BLS-Stx2B were changed to PBS 1X using Amicon 10 kDa MWCO ultra centrifugal filters (Millipore) and concentrated to 1.4 mg/mL. Then, 3 µL of sample were applied onto 1.2/1.3 300-mesh UltrAuFoil grids (Electron Microscopy Sciences). The grids were previously plasma cleaned using a Tergeo-EM apparatus (Pie Scientific LLC). Finally, the loaded grids were blotted for 3 s and plunge-frozen in nitrogen cooled liquid ethane using an EM GP2 machine (Leica).

For the preparation of the pFab-bound complexes, samples of BLS-Stx1B and BLS-Stx2B chimeras were mixed with the corresponding pFabs in 1.3X molar excess and incubated for 1 h at room temperature. Then, the mixture was subjected to SEC chromatography to separate complexes from unbound pFabs. Elution was carried out with PBS 1X and 1 M urea on a Superdex200 increase column (Cytiva) on an Akta Pure System (GE Healthcare). The fractions containing complexes were buffer exchanged to PBS 1X and concentrated to 0.6 mg/mL using Amicon 10 kDa MWCO ultra centrifugal filters (Millipore). The samples were immediately used for cryo-EM sample preparation. Carbon C-flat R1.2/1.3 300-mesh grids (Protochips Inc.) were coated with 30 nm gold using a Leica EM ACE600 Sputter Coater and plasma cleaned using a Tergeo-EM apparatus (Pie Scientific LLC). Then, 3 µL of sample were applied onto the grids, blotted for 3 s and plunge-frozen in liquid ethane using an EM GP2 (Leica) machine.

### Cryo-EM data collection, image processing and model building

In all cases movies were recorded on a FEI Titan Krios electron microscope (Thermo Fisher Scientific) at 300 kV equipped with a K3 direct electron detector (Gatan). Data collection parameters are listed in Table 1.

The image processing of BLS-Stx1B and BLS-Stx2B was analogous, and a detailed description can be found in Fig. S1 and S2, respectively. Briefly, the raw movies were motion corrected using MotionCor2 [43]. Further image processing was performed in CryoSPARC v4.1 [36]. The contrast transfer functions (CTF) were calculated by Patch CTF using the aligned and dose-weighted images. 2D templates were generated by first picking particles with blob picker, extracting and 2D classifying particles. These templates were used to pick particles which were then extracted and binned by 4. The resulting particles were interpolated in a subset of good micrographs using Curate exposures job to obtain a set of good particles and to train an AI picking model using Topaz [37]. This model was used further in a Topaz extraction job. The resulting particles were extracted, binned by 4 and combined with the previous set of particles. After removing duplicates, the particles were re-extracted without binning. Ab-initio reconstructions were performed with these particles. Further refinements were performed using homogeneous refinement with D5 symmetry imposed. The resulting 3D-aligned particles were subjected to local CTF refinement and then used again for homogeneous refinement. Local resolution estimation and filtering were performed in CryoSPARC v4.1 [36]. The final maps were sharpened with AutoSharpen of Phenix [44,45]. The consensus volumes and the corresponding particles were used in a 3D Flexibility [31] pipeline to assess the movement of the toxin subunits. In both cases, meshes were constructed over the obtained consensus volumes and the motion model was trained with four latent coordinates. The movements were assessed by reconstructing multi-frame volume series from the latent space and rendered for visualization in UCSF Chimera [46].

Crystallographic models of Stx1B (PDB code: 1QNU) [15], Stx2B (PDB code: 3MXG) [47] and BLS (PDB code: 1XN1) [29] were fitted into the cryo-EM maps using fit-in-map in ChimeraX [48]. Manual building in COOT [49] and real space refinement in Phenix were performed iteratively. Molprobity [50] was used to validate the model and the corresponding statistics are reported in Table 1. Figures and movies were prepared with UCSF Chimera [46] and ChimeraX [48].

For the immune complexes of both BLS-Stx1B and BLS-Stx2B the image processing was analogous, and a detailed description can be found in Fig. S4 and S5, respectively. Initial processing was performed as described for the unbound chimeras. Particles corresponding to the immune complexes were picked using Topaz with pre-trained models from the corresponding unbound chimera data. After extracting and 2D classifying the particles, Ab-initio reconstructions were performed. The resulting volumes were used as references for homogeneous refinement without imposed symmetry. A 20 Å low-pass filtered volume of the BLS core region and a mask covering this part were used in both cases to run a second 3D reconstruction applying D5 symmetry. Afterwards, symmetry expansion over D5 was performed and the resulting particles were subjected to 3D classification. First, focused classification was performed, using spherical masks covering the extra density accounting for each bound Fab. A second 3D classification round was performed on each class showing clear Fab density without focused mask. The resulting maps of each individualized complex were denoised by low-pass filtering to unmasked resolution in CryoSPARC v4.1 [36].

## Acknowledgments

This work was supported by the Argentinean Ministry of Science and Technology (MINCyT), the Argentinean Research Council (CONICET), and the National Agency for the Promotion of Science and Technology of Argentina (ANPCyT), under grant PICT 2020-3047. Part of this research was supported by the US National Institute Health Intramural Research Program; US National Institutes of Environmental Health Sciences (ZIC ES 103326 to MJB) and the NIH Intramural Targeted Anti-COVID-19 Program (ZIA ES 103341 to MJB). We are grateful to all members of the Molecular Microscopy Group from GISBL, NIEHS for their additional help on cryo-EM sample preparation and data collection. MLC, FAG, VZ, and LHO are researchers from CONICET. AEC and LHO gratefully acknowledge CONICET for providing fellowship and travel support to access cryo-EM facilities for data acquisition.

## Accession Numbers

Cryo-EM maps of the chimeric proteins were deposited in the Electron Microscopy Data Bank (http://www.ebi.ac.uk/pdbe/emdb/) under the accession codes EMD-42072 (BLS-Stx1B) and EMD-42073 (BLS-Stx2B). The associated atomic models were deposited into the Protein Data Bank with accession codes 8UAV and 8UAW, respectively. The cryo-EM maps of the chimeras complexed with the pFabs were deposited in the Electron Microscopy Data Bank (http://www.ebi.ac.uk/pdbe/emdb/) under the accession codes EMD-48123, EMD-48124, EMD-48125, EMD-48126, EMD-48127, EMD-48128, EMD-48129, EMD-48130.

