## Supplementary Materials for "Cryo-EM structures of engineered Shiga toxin-based immunogens capable of eliciting neutralizing antibodies with therapeutic potential against Hemolytic Uremic Syndrome"

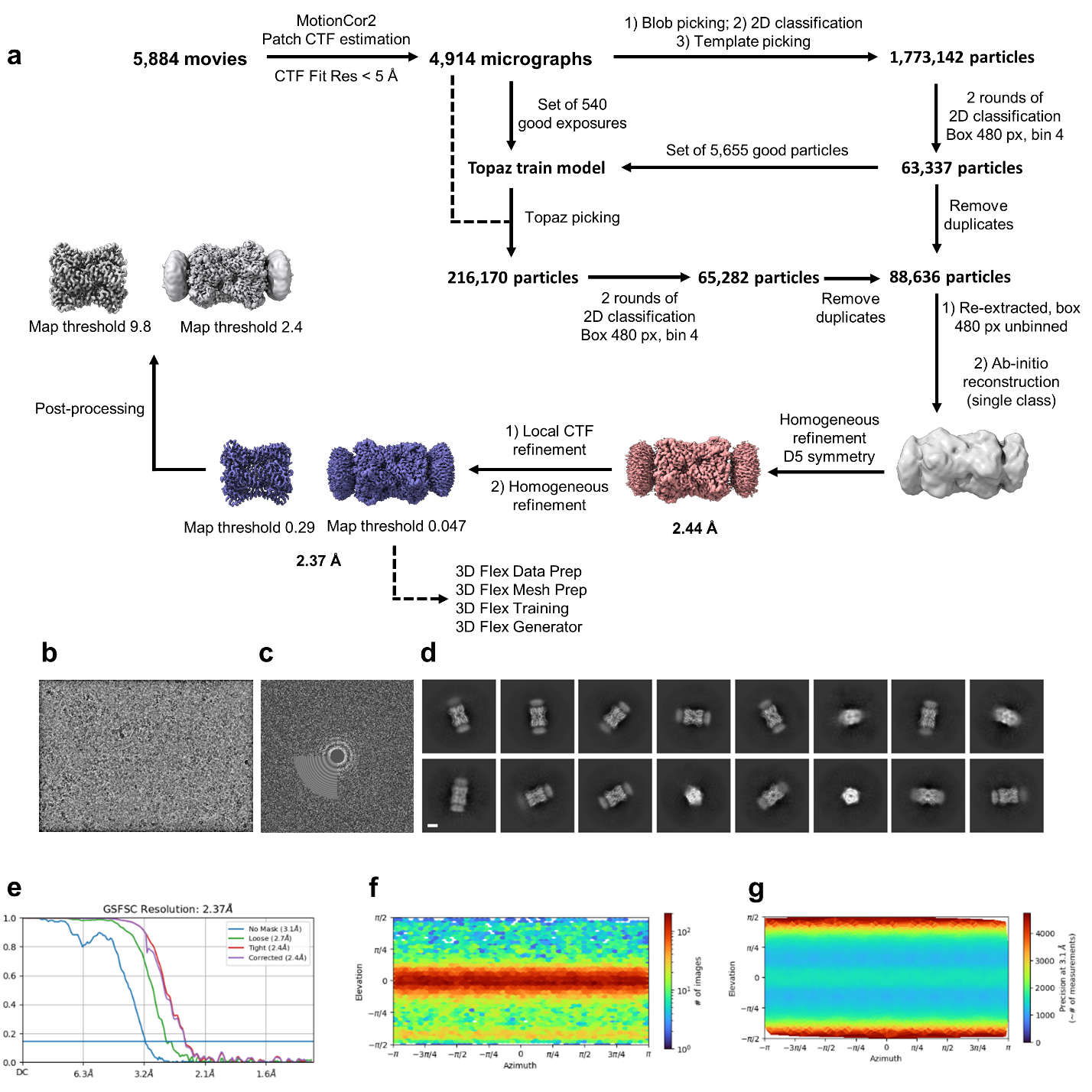

**Figure S1. Cryo-EM data processing of BLS-Stx1B chimera. a.** Data processing workflow. **b.** Representative motion-corrected cryo-electron micrograph. **c.** Corresponding 2D power spectrum. **d.** Reference-free 2D class averages with a 50 Å scale bar. **e.** Gold-standard Fourier shell correlation (FSC) curves for BLS-Stx1B 3D reconstruction. **f.** Angular distribution of particles. **g.** Posterior precision directional distribution of particles.

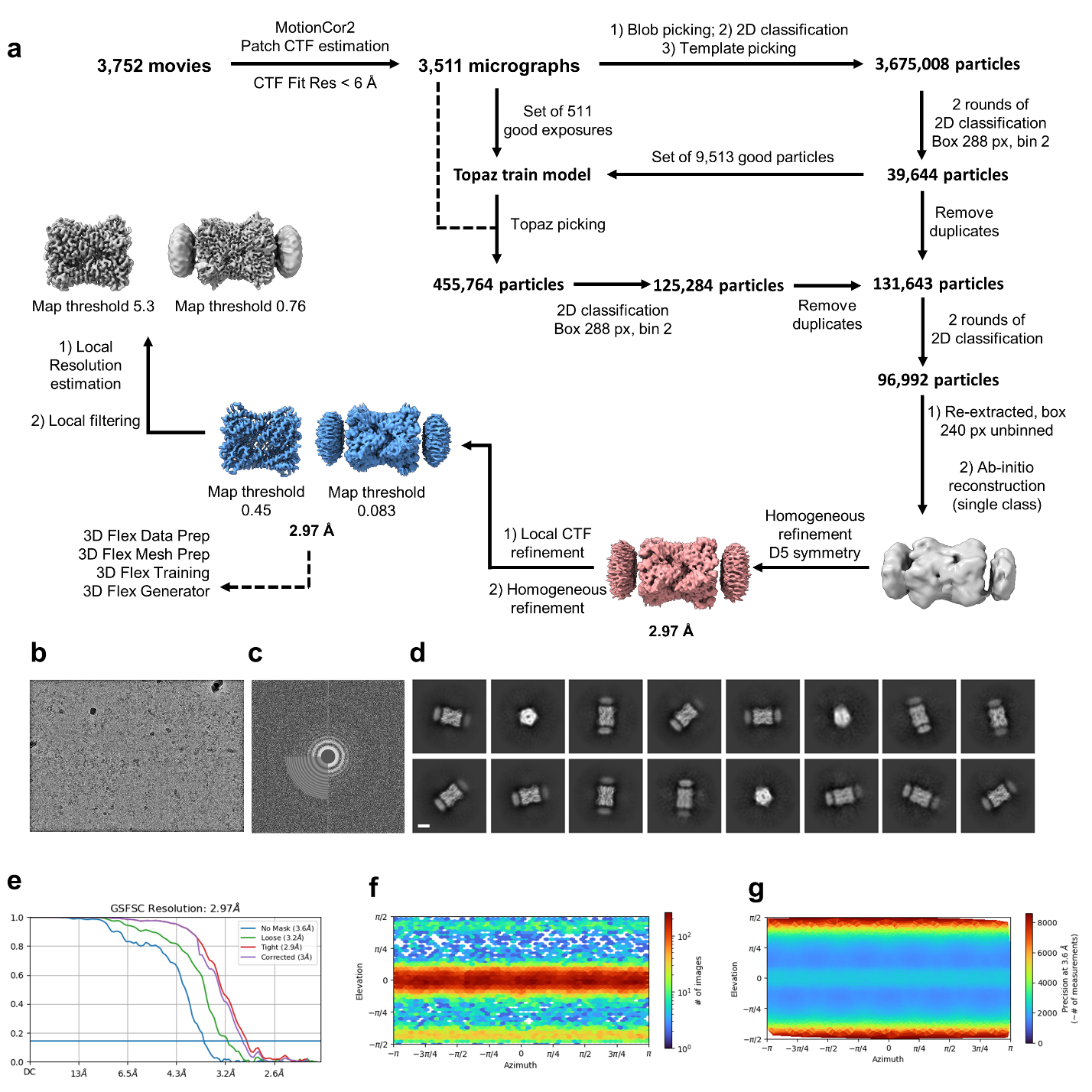

**Figure S2. Cryo-EM data processing of BLS-Stx2B chimera. a.** Data processing workflow. **b.** Representative motion-corrected cryo-electron micrograph. **c.** Corresponding 2D power spectrum. **d.** Reference-free 2D class averages with a 50 Å scale bar. **e.** Gold-standard Fourier shell correlation (FSC) curves for BLS-Stx2B 3D reconstruction. **f.** Angular distribution of particles. **g.** Posterior precision directional distribution of particles.

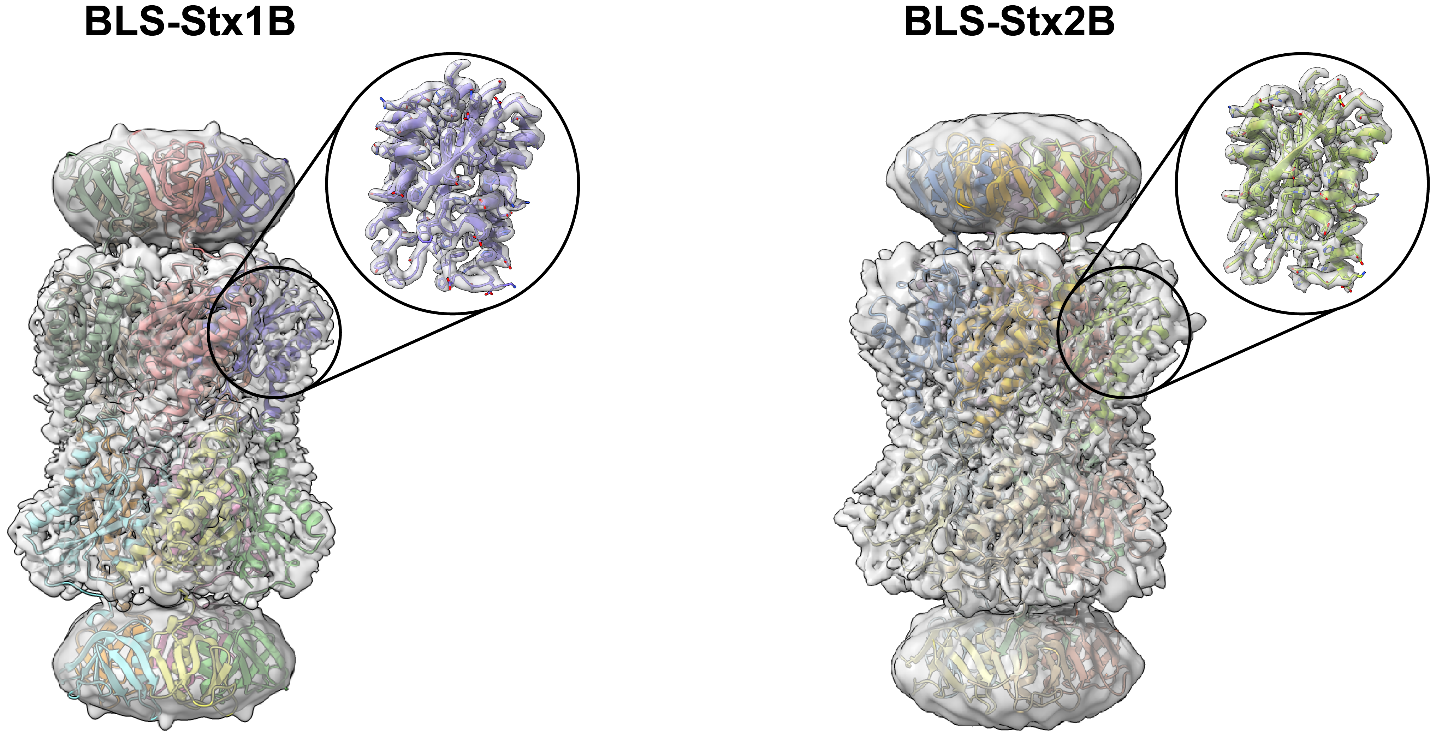

**Figure S3. Atomic models BLS-Stx1B (*left*) and BLS-Stx2B (*right*) chimeras fitted in the respective cryo-EM maps.** The models are shown in ribbon representation, colored by chains and displayed with the maps (gray) transparently overlaid. The insets show a segment of the model corresponding to a BLS monomer.

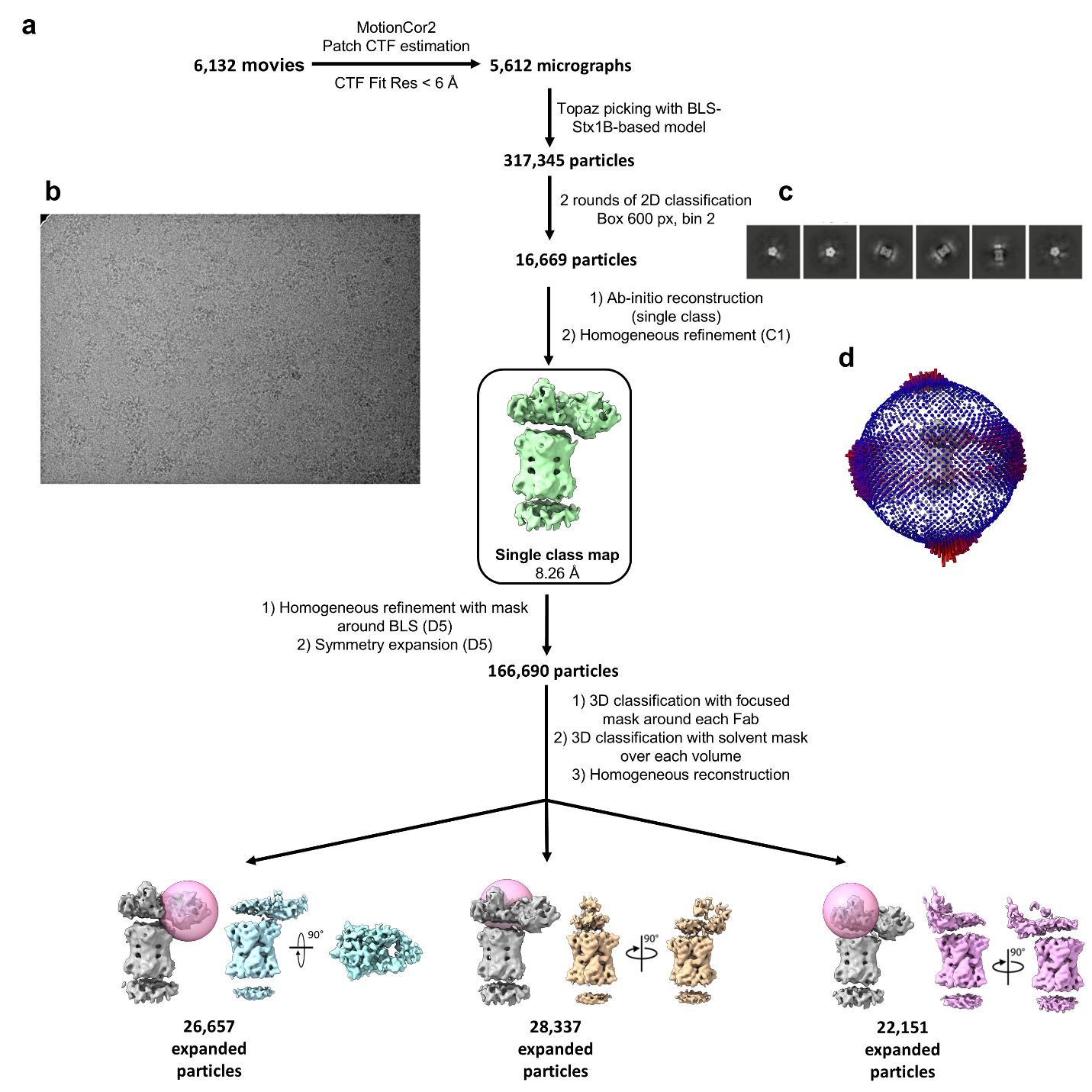
**Figure S4. Cryo-EM data processing for BLS-Stx1B complexed with pFabs. a.** Data processing workflow using Topaz picking. Inset square remarks the single class map obtained for the BLS-Stx1B/pFabs complexes. Pink sphere represents the focused mask used during focused 3D classification. **b**. Representative micrograph. **c.** 2D class averages. **d.** Angular distribution 3D plot.

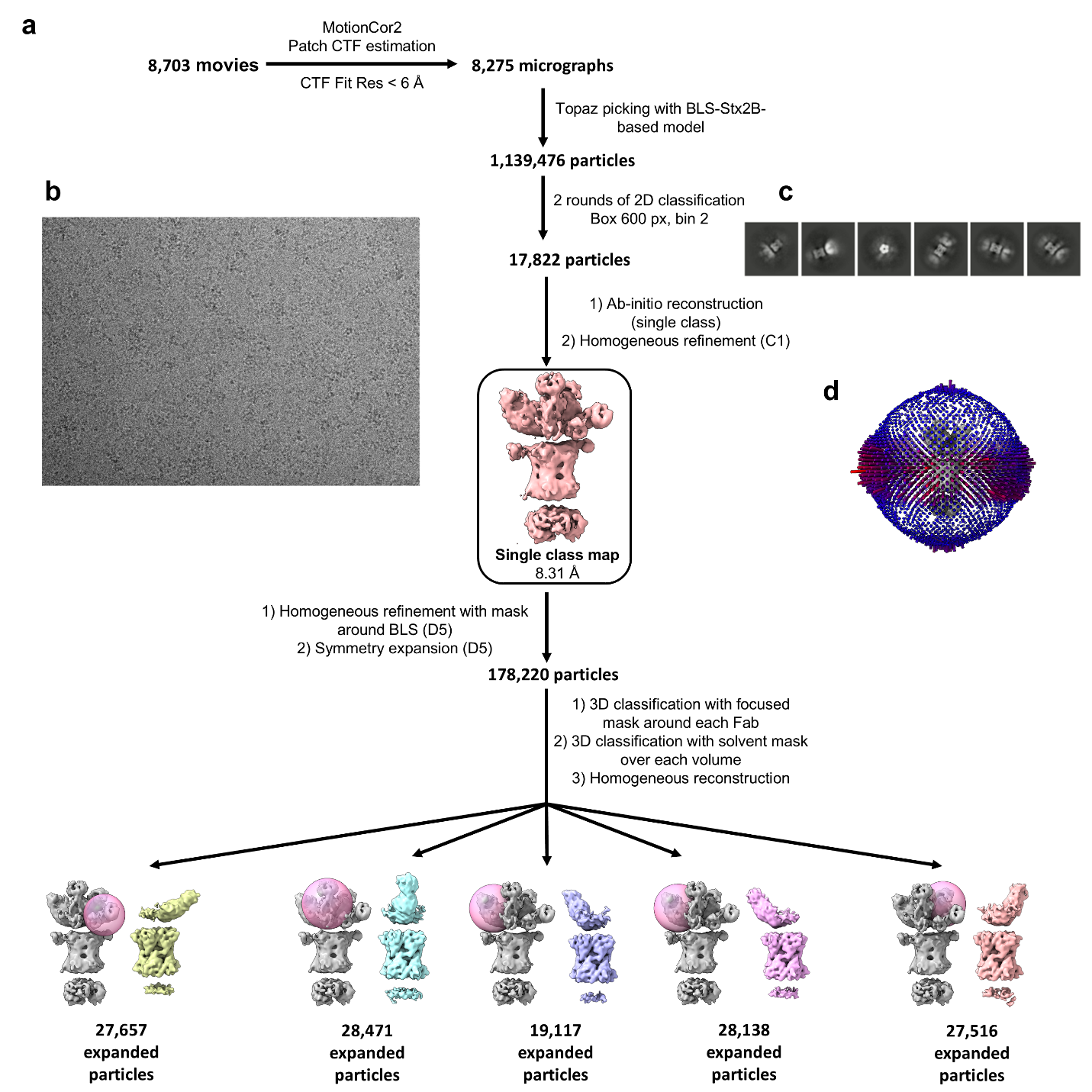
**Figure S5. Cryo-EM data processing for BLS-Stx2B complexed with pFabs. a.** Data processing workflow using Topaz picking. Inset square remarks the single class map obtained for the BLS-Stx2B/pFabs complexes. Pink sphere represents the focused mask used during focused 3D classification. **b**. Representative micrograph. **c.** 2D class averages. **d.** Angular distribution 3D plot.
